# Microbial-derived D-lactate and LPS shape growth and inflammatory signalling in endometrial glandular epithelium

**DOI:** 10.64898/2026.03.09.710619

**Authors:** Lucía Blanco-Rodríguez, Apostol Apostolov, Amruta Pathare, Darja Lavõgina, Merli Saare, Reet Mändar, Signe Altmäe, Andres Salumets, Alberto Sola-Leyva

**Author notes:** Correspondence to: Alberto Sola-Leyva,; Andres Salumets,; Signe Altmäe.

## Abstract

The endometrium, the inner lining of the uterus, is a dynamic tissue that undergoes precise molecular and structural changes to achieve a receptive state capable of supporting embryo implantation. Although the uterine environment was long considered sterile, molecular studies have detected microbial signals and bioactive compounds that may influence endometrial function. Endometrial epithelial organoids (EEOs) provide a three-dimensional in vitro model that recapitulates the architecture, polarity, and hormonal responsiveness of native endometrial tissue. This study aimed to elucidate how bacterial-derived compounds, including D-lactate (D-lac), commonly associated with *Lactobacillus* communities, and lipopolysaccharides (LPS), a component of Gram-negative bacteria, affect the transcriptomic profile of the endometrial epithelium under a hormonally induced receptive state. EEOs were exposed to different concentrations of these compounds, and relative metabolic activity was monitored through resazurin-based assays, revealing no significant alterations across the conditions tested. Transcriptomics analysis of hormonally stimulated EEOs, mimicking the mid-secretory phase, revealed that D-lac modulated genes related to epithelial development, tissue remodelling and growth regulation, whereas LPS influenced genes associated with inflammatory signalling and immune response. While key markers of receptivity remained largely stable, small transcriptional changes suggest that microbial signals may modulate the functional balance of the receptive endometrium. These findings highlight a modulatory role of microbial signals on endometrial epithelial function and demonstrate that EEOs are a robust platform for exploring host–microbe interactions in the uterus, offering new insights into the mechanisms underlying uterine receptivity.

## Introduction

Successful embryo implantation requires the acquisition of endometrial receptivity, a transient physiological state in which the endometrium undergoes molecular and structural remodelling that enables blastocyst adhesion and invasion [1]. Traditionally, the female upper reproductive tract was considered sterile, a view largely based on the limitations of early microbial detection techniques. However, advances in sequencing technologies have revealed microbial signals in the uterus, challenging this long-held assumption [2]. Despite these findings, the presence of active microorganisms in the healthy human endometrium remains debated and many questions about their biological relevance are still unresolved [3].

Although it remains unclear whether microorganisms detected in the endometrium represent true residents or transient populations ascending from the vagina, their DNA and molecular products can be identified within the endometrial environment, suggesting the potential for functional effects [4]. Importantly, these microbial signals may influence the molecular pathways required for the acquisition of endometrial receptivity, thereby modulating the endometrium’s readiness for implantation [5]. The specific impact of microbial signals likely depends on the composition of the local or adjacent microbial communities, which determines the types and abundance of bioactive compounds present. A reproductive tract microbiome dominated by *Lactobacillus* species has been consistently associated with improved endometrial receptivity, implantation success and favourable reproductive outcomes, whereas a non-*Lactobacillus*–dominated microbiome, including the presence of potentially pathogenic bacteria such as *Escherichia coli*, has been linked to local inflammation and reduced implantation and pregnancy rates [4, 6].

In the case of a reproductive tract microbiota dominated by *Lactobacillus* species, bacterial metabolites such as lactate are highly enriched. Lactate is a chiral molecule that exists as two enantiomers forms, L-lactate (L-lac) and D-lactate (D-lac), due to the asymmetry of its central carbon atom. In humans, central metabolic pathways such as glycolysis are highly stereospecific and generate almost exclusively L-lac, reflecting a broader biological preference for L-configured metabolites [7]. This preference applies not only to energy metabolites such as lactate, but also to amino acids, as ribosomes incorporate exclusively L-amino acids [8]. In contrast to human cells, many bacteria possess distinct stereospecific lactate dehydrogenases that enable the synthesis of D-lac, L-lac, or both, highlighting fundamental metabolic differences between human and bacteria in the handling of chiral metabolites. Within the vaginal microbiota, *Lactobacillus* species display characteristic species-specific lactate stereochemistry. *L. crispatus* produces both enantiomers, *L. jensenii* predominantly produces D-lac, whereas *L. iners* produces only L-lac [9, 10]. In the vaginal environment, it is estimated that less than 15% of total lactate derives from anaerobic metabolism of host epithelial cells, indicating that the majority of lactate is produced by the resident microbiota [11, 12]. Because human cells do not generate D-lac, this enantiomer constitutes a specific biomarker of microbial metabolic activity. D-lac has been shown to play a central role in maintaining vaginal mucosal homeostasis by supporting epithelial barrier integrity and limiting pro-inflammatory signalling [9, 13]. Beyond these functions, however, whether D-lac exerts similar regulatory effects on endometrial epithelial cells remains largely unexplored. Investigating its potential to modulate endometrial immune and metabolic pathways is therefore critical for understanding how signals originating from a *Lactobacillus*-rich microbiota may contribute to the establishment of a receptive endometrial state.

Conversely, conditions characterized by enrichment of Gram-negative bacteria are associated with increased levels of others microbial components, particularly lipopolysaccharides (LPS). LPS is a class of structurally complex glycolipid composed of three main domains. Lipid A, which anchors the molecule to the bacterial membrane, consists of a phosphorylated di-glucosamine backbone substituted with multiple acyl chains. Linked to lipid A through the core oligosaccharide, the O antigen extends outward from the bacterial surface and is formed by repeating oligosaccharide units whose structural variability among strains underlies antigenic diversity and shapes host immune recognition [14]. As the most exposed region of the molecule, the O antigen plays a key role in host–microbe interactions, including modulation of complement activation and the induction of strain-specific antibody responses. These molecular features underlie the capacity of LPS derived from Gram-negative species, such as *E. coli,* can activate innate immune pathways, promote chronic inflammation and impair epithelial function and local immune tolerance [15–17].

The receptive endometrium exhibits a distinctive transcriptomic profile, characterized by the regulation of genes involved in epithelial adhesion, barrier function, immune modulation, and tissue remodelling [18, 19]. Given the central role of the endometrial epithelium in mediating interactions between the uterine environment and the implanting embryo, microbial signals mediated by D-lac and LPS may directly influence epithelial cell functions. To study these effects in a controlled and physiologically relevant context, advanced *in vitro* models are essential. Three-dimensional endometrial epithelial organoids (EEOs) faithfully recapitulate the architecture, polarity, and hormonal responsiveness of the native endometrium, making them a powerful platform for functional studies [20, 21]. Their stability and responsiveness to steroid hormones allow for the detailed examination of how microbial signals impact the molecular mechanisms leading endometrial receptivity.

In the present study, EEOs derived from fertile donors were used to explore the transcriptomic and functional effects of bacterial-derived compounds, including D-lac and LPS (Figure 1). Through transcriptomic profiling, we aimed to elucidate how microbial signals regulate endometrial epithelial function during a receptive state.

**Figure 1.**
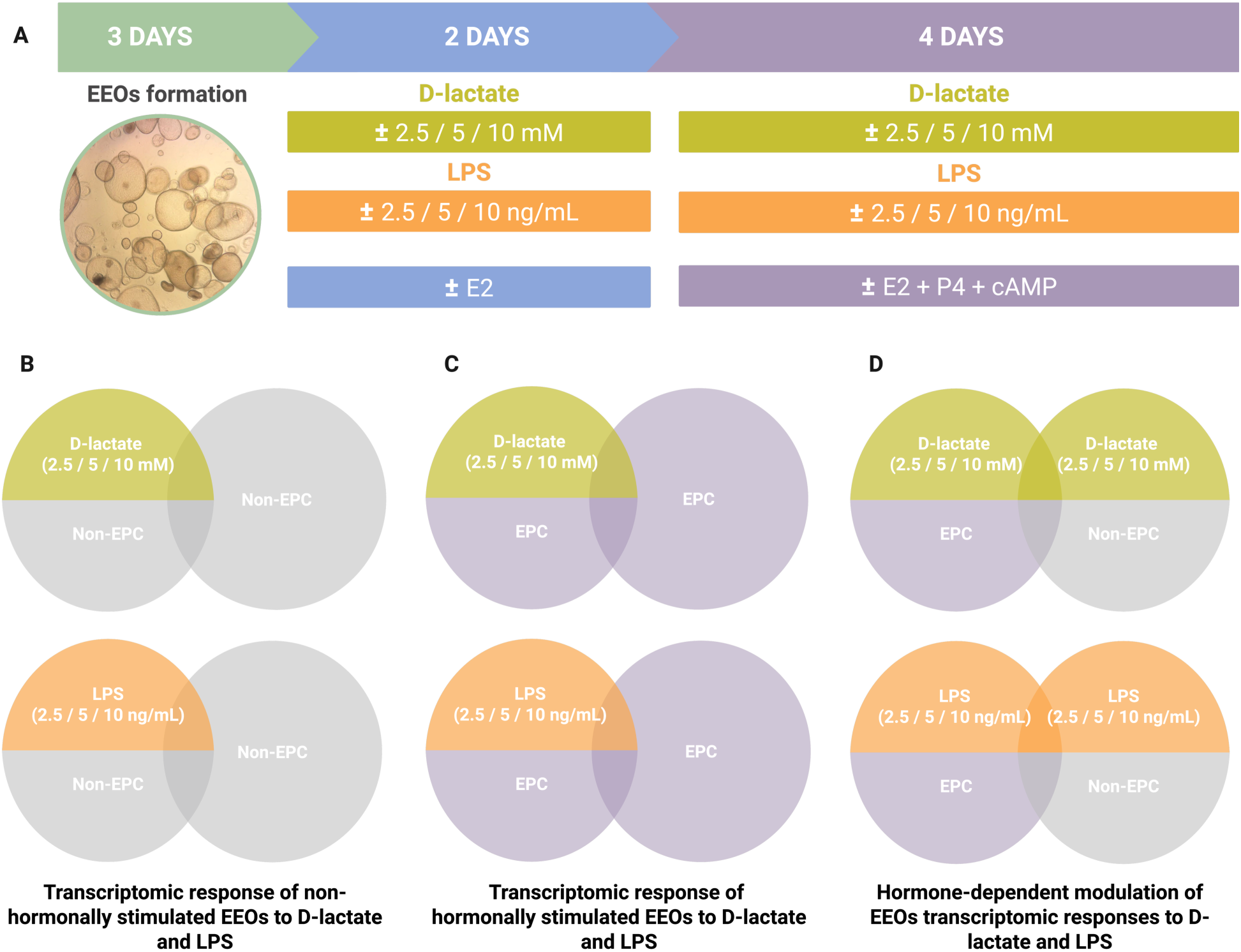
Experimental workflow and transcriptomic analysis strategy. The assay workflow began with endometrial epithelial cell seeding, followed by a 3-day period for organoid formation. Once established, endometrial epithelial organoids (EEOs) were exposed for 2 days to estradiol (E2) in combination with increasing concentrations of D-lactate (2.5 mM, 5 mM, and 10 mM) or lipopolysaccharide (LPS) (2.5 ng/mL, 5 ng/mL, and 10 ng/mL). Organoids were then exposed for 4 days to estradiol (E2), progesterone (P4), and 8-bromo-cyclic adenosine monophosphate (cAMP), while maintaining exposure to D-lactate or LPS at the same concentrations (A). Below, the comparisons performed to identify transcriptomic changes are shown, addressing three main objectives: transcriptomic response of non-hormonally stimulated (Non-EPC) EEOs to D-lactate and LPS (B), transcriptomic response of hormonally stimulated (EPC) EEOs to D-lactate and LPS (C) and hormone-dependent modulation of EEOs transcriptomic responses to D-lactate and LPS (D).

## Materials and methods

### Ethical approval

The study was approved by the Research Ethics Committee of the University of Tartu, Estonia (No. 398/M-7), and written informed consent was obtained from all participants.

### Participants and sample collection

Fertile women of reproductive age were recruited from South Estonian Hospital (Võru, Estonia). The inclusion criteria required participants to have at least one live-born child and to avoid hormonal medications for at least three months prior to the study. Participants were excluded if they had sexually transmitted diseases, uterine pathologies, endometriosis or polycystic ovary syndrome. The phase of the menstrual cycle was determined through menstrual history and detection of the luteinizing hormone (LH) peak using the BabyTime hLH urine test (BR-3645, Pharmanova, Belgrade, Serbia). Endometrial biopsies (N=3) were obtained at two menstrual cycle phases: two from early secretory phase (LH+3, LH+5), and one from late secretory (LH+12). The mean age of the participants was 31.33 ± 3.51 years, and the mean body mass index (BMI) was 23.7 ± 3.97 kg/m². Endometrial biopsies were obtained using a Pipelle flexible suction catheter (Laboratoire CCD, Paris, France) and immediately placed in HypoThermosol FRS Preservation Solution (Sigma, St. Louis, MO, USA). The samples were quickly transported to the laboratory for processing and initiation of organoid culture.

### Establishment of endometrial organoids and single cell dissociation

Epithelial glands were isolated from endometrial biopsies (N = 3) via tissue dissociation, and EEOs were subsequently generated and cultured as previously described [21]. Specifically, the endometrial biopsy was transferred to a sterile tube and washed with RPMI 1640 medium (1X, 11875093, Thermo Fisher Scientific, Waltham, MA, USA) supplemented with 1% penicillin-streptomycin (15140148, Thermo Fisher Scientific) and incubated at 37 °C with agitation until no visible blood residues remained. The tissue was then minced with a scalpel and enzymatically dissociated using a combination of Dispase II (10 mg/mL, D4693, Sigma-Aldrich) and collagenase V (2 mg/mL, C9263, Sigma-Aldrich) for 20 minutes, under gentle agitation with vigorous shaking every 5 minutes. Digestion was halted by adding RPMI 1640 medium supplemented with 10% fetal bovine serum (FBS) (A5256701, Thermo Fisher Scientific).

The resulting cell suspension was filtered through a 100 µm sieve (22363549, Thermo Fisher Scientific) to retain epithelial glandular fragments. To recover the fragments, the filter was inverted and washed with RPMI 1640 medium supplemented with 10% FBS, transferring the contents to a new tube. Following centrifugation at 600 *g* for 6 minutes, the pellet was resuspended in Advanced DMEM/F-12 (1X, 12634010, Thermo Fisher Scientific) supplemented with 0.1% Y-27632 (10 µM, 688000, Sigma-Aldrich). After a second centrifugation, the supernatant was discarded, and the pellet was embedded in 100% Matrigel (CLS356231, Corning®, Corning, NY, USA).

Twenty-five-microliter drops of the cell suspension were seeded into 48-well culture plates (677180, Greiner Bio-One, Kremsmünster, Austria) and incubated at 37 °C in a humidified atmosphere with 5% CO₂ in an inverted position for 10 minutes, allowing the Matrigel to solidify and the cells to distribute evenly. After solidification, the plates were returned to the normal orientation, and 250 µL of expansion medium was added to each well and renewed every 48 hours [21]. EEOs were mechanically dissociated and reseeded every 10–14 days. For all experiments, EEOs between passages 2 and 10 were used. To seed a defined number of cells per well (as described in subsequent sections), organoids were enzymatically dissociated into single cells using Dispase II (10 mg/mL).

### Immunostaining

The EEOs were treated with Cell Recovery Solution (Corning) for 30 min to dissolve the Matrigel and subsequently rinsed with cold Advanced DMEM (Thermo Fisher). Samples were fixed in 4% PFA for 45 min, followed by three washes with PBT (0.01% Tween-20 in 1× PBS). Blocking was carried out for 15 min at 4 °C using a solution containing 10% donkey serum, 0.1% Triton X-100, and 0.2% BSA in 1× PBS. Organoids were incubated overnight at 4 °C with the primary antibodies Ki-67 (ab15580, 1:100) and pan-cytokeratin (ab86734, 1:100), diluted in washing buffer (0.1% Triton X-100 and 0.2% BSA in 1× PBS). Following three washing steps, samples were incubated with secondary antibodies Alexa Fluor 647 (anti-mouse, 1:1000, A31571) and Alexa Fluor 555 (anti-rabbit, 1:1000, A31572) for 1 h at room temperature. The EEOs were then washed three additional times and incubated with Hoechst 33342 (H3570, 1:1000) for 5 min, followed by three further washes with washing buffer. Imaging was performed using a Nikon Ti2 spinning-disk confocal microscope.

### Relative metabolic activity assay

To evaluate the effects of D-lac and LPS on the metabolic activity of EEOs, organoids were exposed to varying concentrations of these compounds, and their relative metabolic activity was subsequently assessed using a resazurin assay, as previously described [22]. Briefly, EEOs were dissociated into single cells and 4000 cells per well were seeded onto mini ring-shaped in 96-well Flat Clear Bottom White Polystyrene TC-treated microplates (Corning). The mini-ring configuration facilitates high-throughput compound testing [23]. After Matrigel solidification, 100 µL of expansion media were added to each well and cells were maintained for three days to allow organoid formation.

EEOs were subsequently treated for six days with expansion media containing D-lac at concentrations of 2.5, 5, or 10 mM (71716, Sigma-Aldrich), or with expansion media containing LPS from *E. coli* serotype O111:B4 at concentrations of 2.5, 5, or 10 ng/mL (L2630, Sigma-Aldrich) according to previously reported concentration ranges used under culture conditions [24–27]. Following the exposure period, the culture medium was removed, and the cells were rinsed with DPBS (Capricorn Scientific, Ebsdorfergrund, Germany). Metabolic activity was then assessed by adding 50 µM resazurin (Sigma) in DPBS supplemented with Ca²⁺ and Mg²⁺ (Capricorn). The plates were placed in a Cytation 5 multi-mode reader (BioTek) and absorbance was recorded at 570 nm and 600 nm (monochromator mode; kinetic readings every 15 min for 2 h; read height 8.5 mm, with lid). Absorbance measurements reflect the ratio between the amount of light absorbed at the wavelengths corresponding to resorufin and resazurin maxima, whereas fluorescence emission, recorded simultaneously, reflects the light emitted by resorufin after excitation. For data analysis, the absorbance ratio (570 nm/600 nm) was calculated for each well and time point and fluorescence emission values were also recorded, both data were analysed independently. Data from negative controls (cells cultured without any of the bacterial-derived compound) were pooled and plotted over time to establish the linear range of the assay. Within this range, measurements corresponding to the same compound, concentration and EEOs line were grouped. The data were then normalized to express relative metabolic activity, assigning 100% to the mean value of control wells and 0% to wells containing only the resazurin solution (blank). Statistical analyses were performed using GraphPad Prism 8.0 (La Jolla, CA, USA). Statistical differences between experimental groups and controls were analysed using one-way ANOVA followed by Dunnett’s post hoc test for multiple comparisons. Adjusted p-values < 0.05 were considered statistically significant. Data are presented as mean ± standard error of the mean (SEM).

### Transcriptomics effects of hormonal stimulation and exposure to D-lac or LPS on EEOs

To recapitulate the molecular profile of receptive endometrium and to assess the combined effects of hormonal stimulation and bacterial compounds on EEOs, organoids were first dissociated into single cells, and 15000 cells were seeded in 25 µL Matrigel droplets in 48-well culture plates. Subsequently, 250 µL of expansion medium were added to each well and the cultures were maintained for three days with medium renewal every 48 hours to allow organoid development. On day 3 after seeding and with EEOs formed, the exposure phase was initiated with a final volume of 250 µL per well. Organoids were cultured in: (i) medium containing D-lac (2.5, 5, or 10 mM) plus 10 nM E2 (E8875, Sigma-Aldrich); (ii) medium containing LPS (2.5, 5, or 10 ng/mL) plus 10 nM E2; (iii) medium containing D-lac or LPS at the same concentrations without hormones; (iv) medium containing only 10 nM E2; or (v) expansion medium alone as an untreated control. On days 6 and 8 after seeding, the medium was replaced using the same treatment conditions. For hormone-containing groups, 10 nM E2, 1 µM P4 (P7556, Sigma-Aldrich), and 1 µM 8-Br-cAMP (1140, Bio-Techne) were added (EPC). Parallel cultures treated with D-lac or LPS alone, hormone-only controls, and untreated controls were also maintained. On day 10 of culture, organoids were collected for RNA extraction (Figure 1A). Each condition was tested in triplicate.

### RNA extraction, sequencing library preparation, and sequencing

Transcriptomic changes induced by the hormonal treatment and the exposure of D-lac or LPS in EEOs were assessed by bulk RNA sequencing. After organoid collection, total RNA was extracted using the RNeasy Micro Kit (74004, Qiagen, Hilden, Germany). The concentration and purity of the extracted RNA were with NanoDrop 2000 spectrophotometer (Thermo Fisher Scientific, USA). RNA samples were stored at –80 °C until further processing. For library preparation, mRNA was enriched through poly(A) selection using the TruSeq Stranded mRNA Library Prep Kit (Illumina, San Diego, CA, USA). The resulting cDNA concentration and fragment size distribution were assessed using the TapeStation 2100 system (Agilent, Santa Clara, CA, USA) with High Sensitivity D1000 ScreenTape. Libraries were sequenced on a NextSeq 1000 system (Illumina) using a single-end 80 bp read configuration.

### Differential expression analysis

Fastq files underwent quality control check (QC) and trimming (Trim Galore) and subsequently aligned to the human reference genome (GRCh38) using HISAT2 within the Galaxy platform [28]. Differential gene expression (DEGs) analysis was performed with DESeq2, applying thresholds of |log₂FC| ≥ 2 and an adjusted p-value < 0.05. Heatmaps were generated using Morpheus (https://software.broadinstitute.org/morpheus), principal component analysis (PCA) plots were created with ClustVis, [29] volcano plots were produced using SRplot [30] and gene ontology (GO) enrichment analysis was conducted with ShinyGO [31].

To assess the effect of bacterial-derived compounds on the expression of endometrial receptivity markers in EEOs under hormonal stimulation, a specific subset of genes was analysed based on previously validated markers of endometrial receptivity. These markers were obtained from the beREADY endometrial receptivity model, [19] and their differential expression was evaluated to identify treatment-associated transcriptional changes. Genes were considered significantly altered when they showed an adjusted p-value < 0.05 and a |log₂FC| ≥ 1.

## Results

### EEOs immunofluorescence and relative metabolic activity assay

The basal expression of epithelial and proliferation markers in EEOs was evaluated by immunofluorescence staining for KRT, confirming their epithelial identity, and Ki-67, indicating active cell proliferation (Figure 2A).

**Figure 2.**
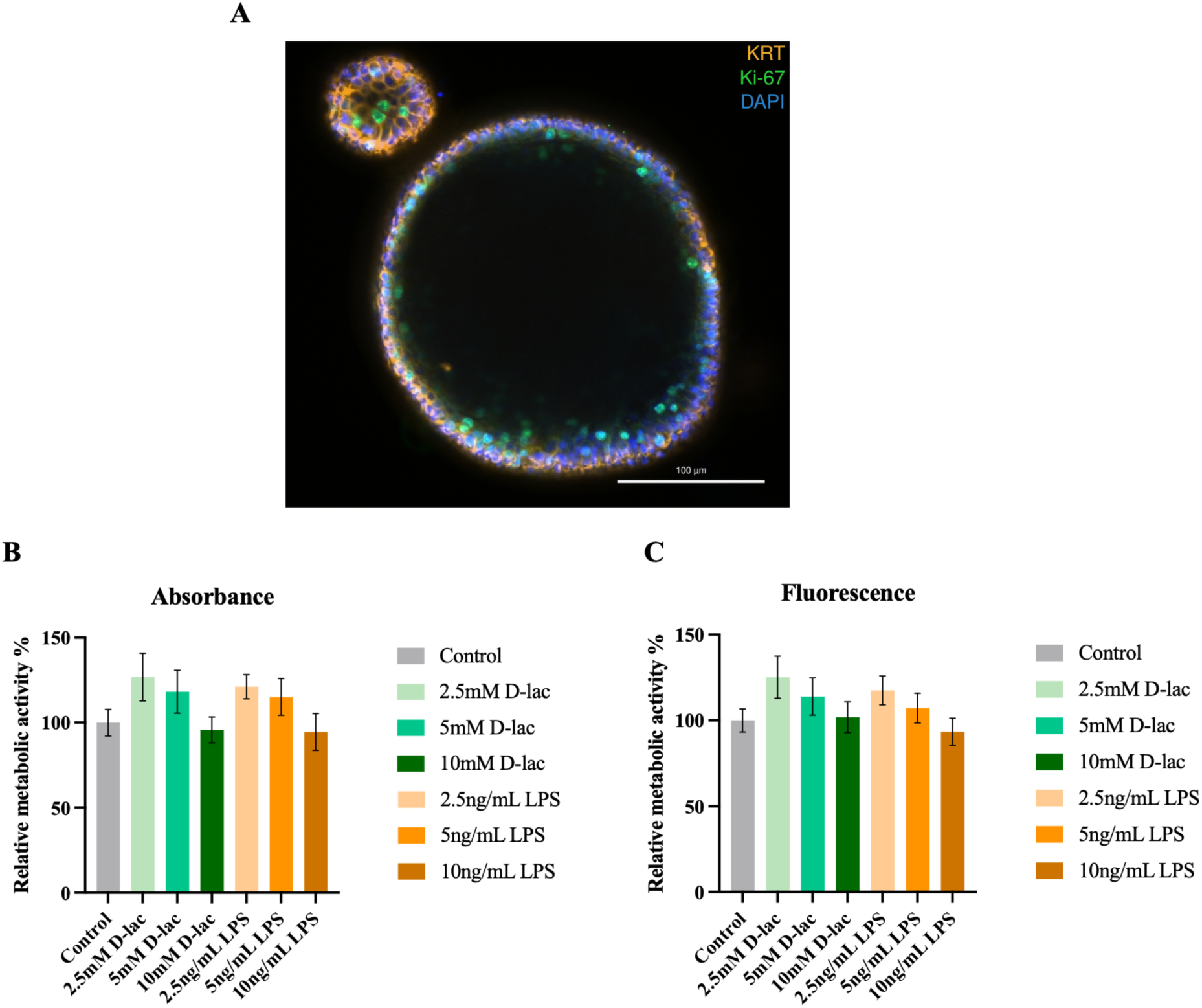
Characterization of endometrial epithelial organoids (EEOs) and their metabolic response to D-lactate (D-lac) and lipopolysaccharide (LPS). EEOs stained for KRT (orange) and Ki-67 (green) (A). Relative metabolic activity was assessed after exposure to increasing concentrations of D-lac and LPS using (B) absorbance measurements at the 570/600 nm ratio and (C) fluorescence intensity measurements. Data are presented as mean ± standard error of the mean (SEM) (N=3). Control group indicates cells cultured without D-lac or LPS. Statistical significance was evaluated using a one-way analysis of variance (ANOVA) followed by Dunnett’s post hoc test for multiple comparisons. No significant differences were observed.

Changes in the absorbance ratio (570nm/600nm) and in fluorescence intensity were monitored during a 2-hour (120 min) incubation of EEOs with resazurin. Based on the control measurements, the linear range of the assay was determined and data corresponding to the last time point within this range were selected for analysis (120 minutes for absorbance and 105 minutes for fluorescence measurements). The effects of D-lac and LPS on the relative metabolic activity of EEOs were evaluated using normalized absorbance and fluorescence data for each experimental condition. Our results demonstrated that D-lac or LPS exposure did not significantly affect the relative metabolic activity of EEOs after 6 days of exposure (Figure 2B-2C).

### Differential gene expression in response to hormonal stimulation

Administration of E2 combined with P4 and 8-Br-cAMP (EPC), which mimics the mid-secretory phase of the menstrual cycle, resulted in marked transcriptional changes in EEOs. Differential gene expression analysis revealed that EEOs hormonally stimulated with EPC exhibited 289 DEGs compared to non-hormonally stimulated EEOs. Among these, 198 genes were upregulated and 91 were downregulated (Figure 3A, Supplementary Table 1). Principal component analysis showed a clear clustering of samples according to the presence or absence of hormonal stimulation (Figure 3B).

**Figure 3.**
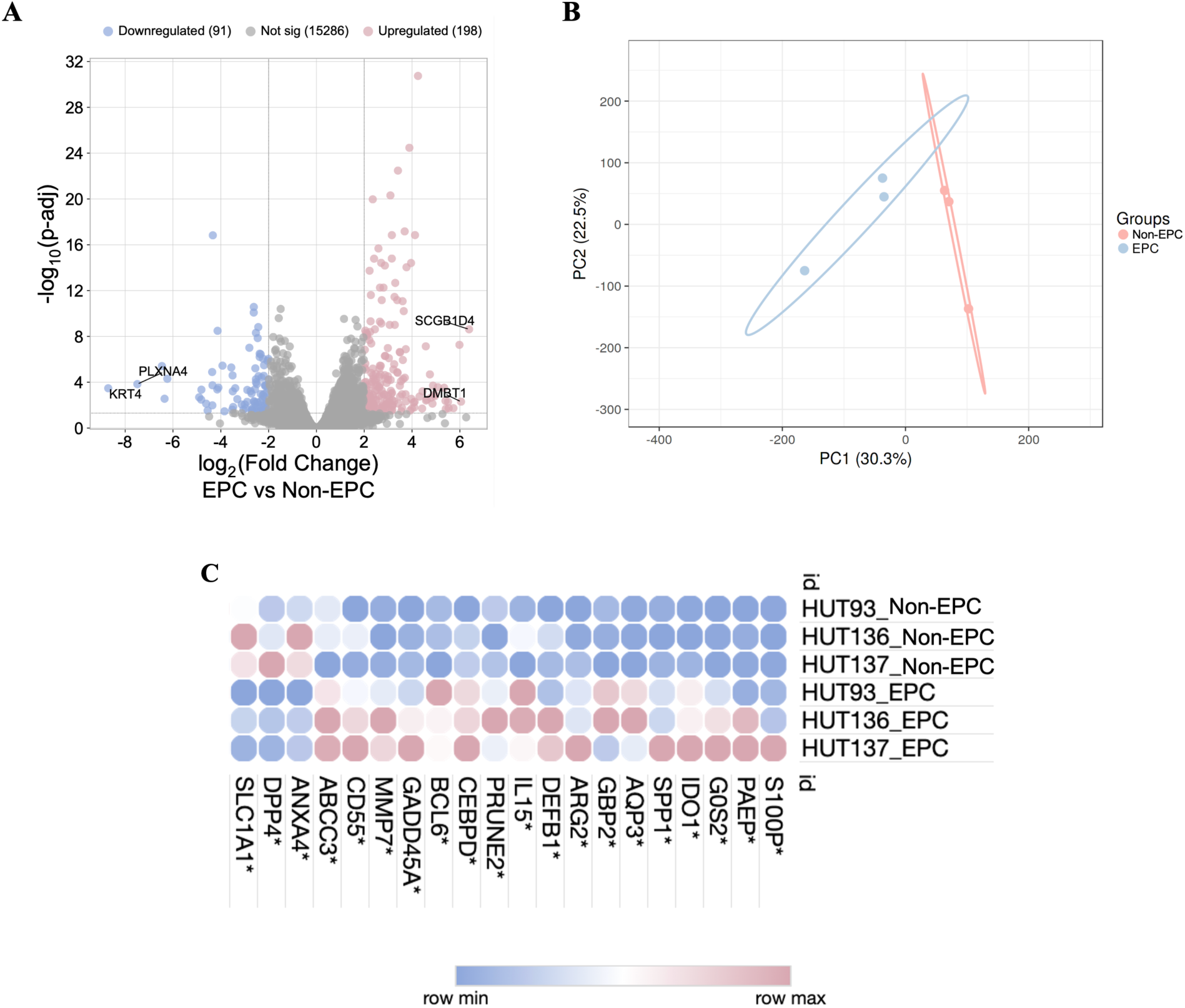
Transcriptomic changes induced by hormonal stimulation in endometrial epithelial organoids (EEOs). A) Volcano plot comparing gene expression profiles between hormonally stimulated EEOs (EPC) and non-stimulated control EEOs (Non-EPC). The plot displays the log₂ fold change (log₂FC) versus the log₁₀ adjusted p-value (p-adj). Differentially expressed genes were defined by an adjusted p-value < 0.05 and |log₂FC| ≥ 2, with upregulated genes shown in pink and downregulated genes in blue. Genes exhibiting the largest positive (*SCGB1D4* and *DMBT1*) and negative (*KRT4* and *PLXNA4*) fold changes are highlighted. B) Principal component analysis (PCA) illustrating the segregation between hormonally stimulated EEOs (EPC) and non-stimulated EEOs (Non-EPC). C) Heatmap illustrating the expression levels of endometrial receptivity markers that are differentially expressed in hormonally stimulated (EPC) EEOs compared to non-stimulated EEOs (Non-EPC). Statistically significant differences are indicated by an asterisk (*), defined as an adjusted p-adj < 0.05 and |log₂FC| ≥ 1.

The EEOs responded robustly to hormonal treatment, displaying substantial modulation of their gene expression profile. Notably, hormonal stimulation led to the upregulation of genes characteristic of the mid-secretory phase, including *PAEP*, *AQP3*, *SPP1*, *LIF*, and *HSD17B2*, indicating the acquisition of a receptive endometrial phenotype. GO analysis revealed significant enrichment of pathways related to regulation of vascular system development, inflammatory response and tissue development in hormonally stimulated EEOs (Supplementary Table 2).

### Identification of markers of endometrial receptivity in hormonally stimulated EEOs

Among the 68 endometrial receptivity markers included in the beREADY model [19] (Supplementary Table 3), 20 were differentially expressed in hormonally stimulated versus non-hormonally stimulated EEOs (Supplementary Table 4). Among these, several key endometrial receptivity-associated genes, including *S100P*, *PAEP*, and *SPP1*, were significantly upregulated (Figure 3C).

### Differential gene expression in response to D-lac or LPS exposure

To identify transcriptomic changes in the EEOs associated with exposure to different concentrations of D-lac or LPS, a differential gene expression analysis was performed. Gene expression profiles of non-hormonally stimulated EEOs exposed to each concentration of D-lac or LPS (non-EPC stimulated + Bacterial-derived compound) were compared with those of non-hormonally stimulated EEOs not exposed to any bacterial-derived compound (non-EPC stimulated) (Figure 1B). This analysis did not reveal significant differences, indicating that, in the absence of hormonal stimulation, none of the compounds tested induced detectable changes in the EEO gene expression profile across the concentrations evaluated.

Secondly, to determine the effect of D-lac and LPS during endometrial receptivity acquisition, we evaluated the gene expression profiles of hormonally stimulated EEOs treated with each concentration of D-lac or LPS (EPC + Bacterial-derived compound) versus those of hormonally stimulated EEOs not exposed to these compounds (EPC) (Figure 1C). No significant differences were observed at any of the LPS concentrations tested. In contrast, treatment with 10 mM D-lac resulted in significant differential expression of *COL1A2*, *MRC2*, and *SMOC1* (adjusted p-value < 0.05 and |log₂FC| ≥ 2) (Supplementary Table 5).

Subsequently, we investigated the impact of hormonal stimulation on the transcriptional responses of EEOs to D-lac or LPS by comparing gene expression profiles of hormonally stimulated EEOs exposed to every concentration of D-lac or LPS (EPC + Bacterial-derived compound) with those non-hormonally stimulated EEOs exposed to the same concentrations (non-EPC stimulated + Bacterial-derived compound) (Figure 1D). Given that hormonal stimulation alone induces substantial transcriptional changes, genes regulated solely by hormonal stimulation were excluded from the analysis. These genes were identified by comparing hormonally stimulated and non-hormonally stimulated EEOs in the absence of bacterial compounds (EPC vs. non-EPC stimulated) (Supplementary Table 1). This strategy enabled the identification of genes whose expression is specifically influenced by the interaction between hormonal signalling and exposure to bacterial compounds. Specifically, 249, 96, and 142 DEGs were identified in the EEOs exposed at 2.5, 5 and 10 mM D-lac, respectively, and 195, 127, and 105 DEGs in the EEOs exposed at 2.5, 5 and 10 ng/mL LPS (Supplementary Table 6). Overall, these results indicate that the transcriptional response of EEOs to D-lac or LPS is dependent on the hormonal context, suggesting that hormonal signalling modulates the ability of EEOs to respond to different bacterial stimuli present in their microenvironment.

To identify biological pathways enriched among differentially upregulated genes under each condition, a GO analysis was performed for each concentration of D-lac and LPS. Based on enrichment analysis, in EEOs exposed to 2.5 mM D-lac, upregulated genes were associated with pathways related to regulation of cell death (*ICAM1*, *TNFAIP3*, *PMAIP1*, *ATF3*) and regulation of cell population proliferation (*SEMA3G*, *TNFRSF1B*, *PLAUR*, *SERPINE2*, *ARG2*, *TNFAIP3*, *ZFP36*, *CCN3*, *DUSP1*, *CD200*) (Supplementary Figure 1A). By contrast, EEOs treated with 5 mM D-lac exhibited no significantly enriched biological pathways. In EEOs exposed to 10 mM D-lac, upregulated genes were associated with pathways involved in growth regulation (*SEMA3G*, *CDA*, *IGFBP5*, *CXCR4*, *MAP1B*, *HEY2*, *SERPINE2*, *IHH*, *AR*, *GJA1*) and epithelial development (*KRT6A*, *MAP1B*, *IGFBP5*, *ID3*, *CXCR4*, *WNT2B*, *HEY2*, *SERPINE2*, *IHH*, *AR*, *IRX3*, *GJA4*, *GJA1*, *CLDN5*, *DHRS9*) (Supplementary Figure 1B).

For LPS exposure, EEOs treated with 2.5 ng/mL showed upregulated genes linked to pathways related to the defence response to symbionts (*OAS3*, *OAS2*, *IFIT3*, *IFIT2*, *RSAD2*, *OASL*, *DDX60*, *HERC5*, *DDX60L*, *BIRC3*) and cytokine response (*LEPR*, *GBP4*, *OAS2*, *ZFP36*, *XAF1*, *AXL*, *CSF1*, *CD38*, *TNFRSF1B*, *IFIT3*, *IFIT2*, *NR1D1*, *OASL*, *TNFRSF13C*, *LAMP3*, *OAS3*, *LAPTM5*, *BIRC3*) (Supplementary Figure 2A). At 5 ng/mL LPS, upregulated genes were associated with pathways related to chronic inflammatory response (*VNN1*, *TNFAIP3*, *IL10*) and positive regulation of cell development (*TNFRSF1B*, *OLFM4*, *SERPINE2*, *CXCR4*, *MAP1B*, *NTRK2*) (Supplementary Figure 2B). Finally, at 10 ng/mL LPS, upregulated genes were linked to the ERBB2 signalling pathway (*AREG*, *HBEGF*, *PTPRR*) and positive regulation of the epidermal growth factor receptor pathway (*AREG*, *HBEGF*, *PLAUR*) (Supplementary Figure 2C).

### Impact of D-lac and LPS on the endometrial receptivity-associated gene expression

To investigate the potential impact of bacterial-derived compounds on endometrial receptivity, differential gene expression was analysed in hormonally stimulated EEOs treated with D-lac or LPS (EPC + Bacterial-derived compound) compared to non-hormonally stimulated EEOs exposed to the same compounds (non-EPC stimulated + Bacterial-derived compound), and focusing on receptivity-associated genes proposed in the beREADY model [19].

Specifically, among the 68 endometrial receptivity markers included in the beREADY model, 17 genes were significantly upregulated and 6 downregulated at 2.5 mM D-lac, 7 significantly upregulated and 5 downregulated at 5 mM D-lac, and 9 significantly upregulated and 6 downregulated at 10 mM D-lac (Figure 4A, Supplementary Table 7). For LPS, 18 genes were significantly upregulated and 2 downregulated at 2.5 ng/mL, 11 significantly upregulated and 5 downregulated at 5 ng/mL, and 13 significantly upregulated and 4 downregulated at 10 ng/mL (Figure 4B, Supplementary Table 7). Notably, *DYNLT3* was uniquely differentially upregulated at 2.5 mM D-lac, *MT1G* uniquely differentially downregulated at 2.5 mM D-lac, *MAP3K5* uniquely differentially upregulated at 2.5 ng/mL LPS, and *TCN1* uniquely differentially downregulated at 10 ng/mL LPS, indicating that these DEGs were not observed in the hormonally stimulated EEOs when compared with non-hormonally stimulated EEOs.

**Figure 4.**
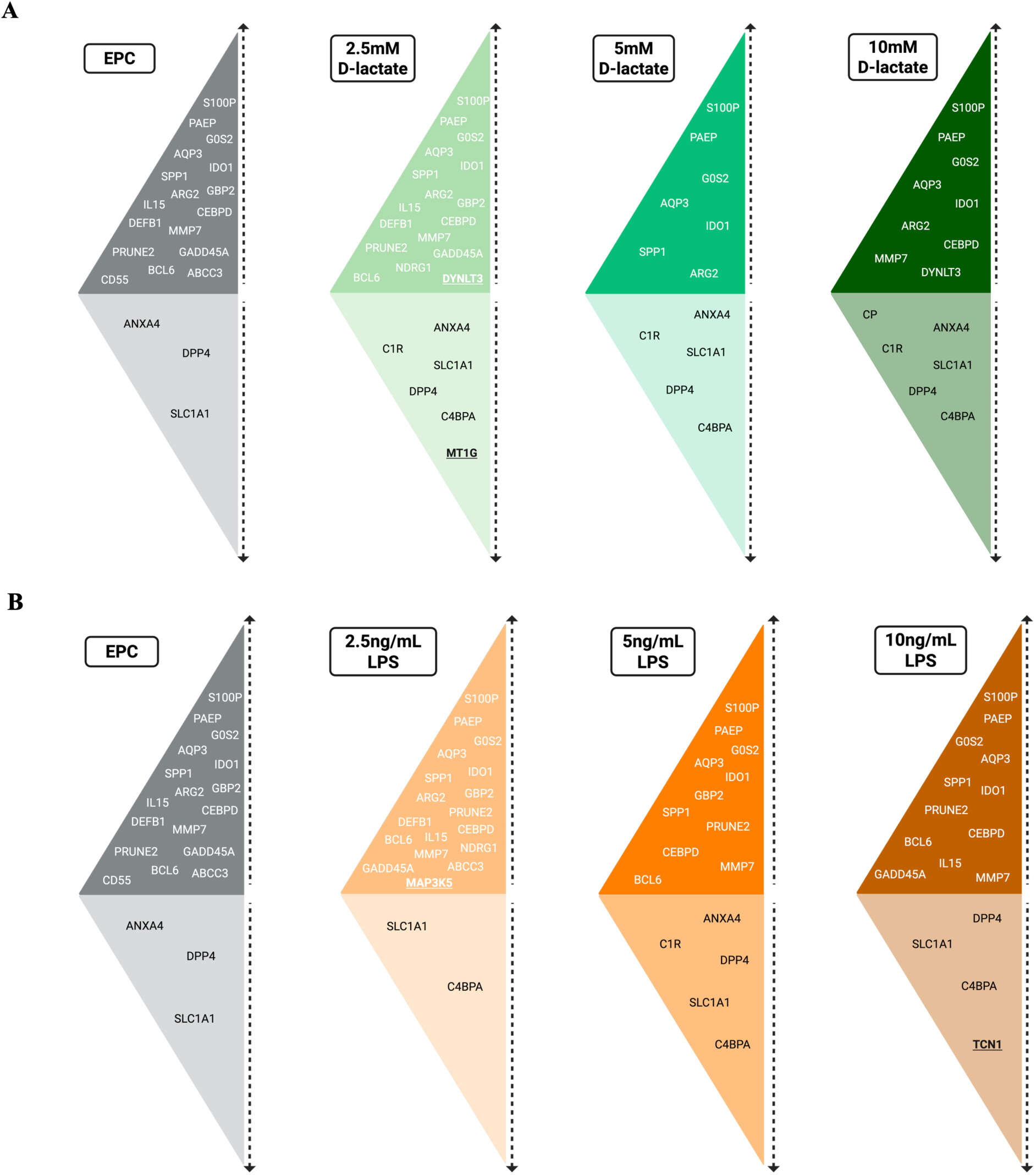
Impact of D-lactate exposure on the expression of endometrial receptivity markers in hormonally stimulated endometrial epithelial organoids (EEOs). Markers significantly upregulated (adjusted *p*-value < 0.05 and log₂ fold change log₂(FC) ≥ 1) or downregulated (adjusted *p*-value < 0.05 and log₂(FC) ≤ −1) in D-lactate–treated EEOs are shown relative to hormonally stimulated EEOs not exposed to D-lactate (EPC). Genes uniquely differentially regulated, indicating that these DEGs were not observed in EPC-EEOs when compared with Non-EPC-EEOs, are underlined.

## Discussion

The endometrial epithelium plays a central role in establishing a receptive environment for embryo implantation by integrating hormonal, immune and environmental signals that collectively support blastocyst attachment and invasion. In addition to host-derived signals, molecules from the surrounding microbial environment may also contribute to modulating epithelial function. EEOs provide a physiologically relevant *in vitro* model to study such interactions. This model has been previously used to investigate host-microorganism interactions, particularly in the context of endometrial pathologies, where exposure to live bacteria has been shown to alter transcriptomic profiles and markers of epithelial function [32]. In the present study, we used EEOs to assess the effects of bacteria-derived compounds rather than direct exposure to live bacteria. Specifically, we focused on D-lac, predominantly produced by a *Lactobacillus*-rich microbiota, and LPS, a marker of increased Gram-negative bacterial abundance, to investigate how these distinct microbial microenvironments influence the transcriptional and functional properties of the endometrial epithelium.

In this study, we focused on D-lac rather than L-lac as a microbial derived compound due to its bacterial origin and its distinct protective roles in the female genital tract [13]. Unlike L-lac, which is produced by both human epithelial cells and bacteria, [33] D-lac is almost exclusively synthesized by certain *Lactobacillus* species, including *L. crispatus*, *L. gasseri*, and *L. jensenii*, but not by *L. iners*, which lack D-lactate dehydrogenase [9, 10]. This selective production is clinically significant, as higher D-lac levels in the vagina correlate with reduced extracellular matrix metalloproteinase inducer (EMMPRIN) expression and matrix metalloproteinase-8 (MMP-8) activity, helping maintain cervical barrier integrity and preventing pathogen ascension to upper reproductive tract [9]. Furthermore, the predominance of D-lac in the vagina reflects a microbiota dominated by protective *Lactobacillus* species, whereas lower D-lac levels, as observed in *L. iners*-dominated or dysbiotic communities, associate with increased susceptibility to bacterial vaginosis and upper genital tract infections [11, 12]. Therefore, D-lac serves both as a functional mediator of epithelial homeostasis and a more specific microbial biomarker than L-lac, providing a clearer link between microbiome composition and reproductive health.

The resazurin assay, which reflects relative metabolic activity, provides an indirect measure of cell viability, as substantial reductions in resorufin formation generally indicate cytotoxic effects of treatment [22]. In our study, none of the tested concentrations of D-lac or LPS caused significant changes compared to the control, suggesting that the treatments were non-cytotoxic. This finding supports the suitability of these experimental conditions for a detailed analysis of transcriptomic alterations, ensuring that cell viability does not confound the interpretation of molecular responses.

The DEGs identified in hormonally stimulated EEOs in the absence of bacterial compounds were consistent with previous reports in EEOs [20, 21, 34] and closely mirrored the transcriptomic profile of the endometrial epithelium *in vivo* during the secretory phase [18]. Notably, among the identified markers, *PAEP* and *SPP1* stand out. *PAEP*, which encodes glycodelin, is positively regulated by progesterone and plays a crucial role in modulating maternal immune tolerance and establishing a supportive environment for embryo implantation [35]. Similarly, *SPP1* (also known as osteopontin), encoding a secreted phosphoprotein involved in embryo adhesion, showed significant upregulation; its expression is specific to the endometrial epithelium and is enhanced during the receptive, mid-secretory phase [18]. These findings reinforce the utility of EEOs as a model for investigating the molecular mechanisms that regulate the implantation window, allowing controlled studies of epithelial-specific responses [21].

In the absence of hormonal stimulation, exposure of EEOs to the bacterial compounds evaluated did not induce significant transcriptional changes, regardless of the concentration tested. This finding suggests that epithelial responsiveness to microbial signals is strongly conditioned by the hormonal environment. The transcriptional response of EEOs to D-lac was detectable only in the presence of hormonal stimulation, highlighting the role of hormones in modulating epithelial sensitivity to microbial-derived signals. Consistent with this, it has been shown that hormonally stimulated EEOs exposed to 10 mM lactate exhibit pronounced alterations in epithelial junctional markers, including decreased ZO-1 and E-cadherin expression alongside increased VIMENTIN levels, closely resembling the features of the secretory-phase endometrium in vivo [24]. These observations suggest that the effects of lactate on epithelial structure are most evident in a hormonally responsive context. However, it is not specified whether D- or L-lac was used in their study, making it unclear which isoform drives the observed effects.

The enrichment of pathways associated with the regulation of cell death and cell population proliferation observed upon treatment with 2.5 mM D-lac in hormonally stimulated EEOs suggests a potential role of D-lac in modulating proliferative control mechanisms. Previous studies using L-lactate demonstrated a direct inhibitory effect on endometrial epithelial cell proliferation. Specifically, exposure of primary endometrial epithelial cells to 5 mM L-lactate was associated with a significant reduction in proliferation–cell index, and treatment of ECC-1 cells with 2.5 mM L-lactate significantly decreased proliferation rates across the culture period (12–48 h) [27]. While our study employed D-lactate and focused on transcriptional responses rather than functional proliferation assays, the enrichment of proliferation-regulatory pathways supports a convergent functional outcome for both lactate isomers in modulating epithelial proliferative control.

In EEOs exposed to 10 mM D-lac, DEGs were enriched in pathways related to growth regulation and epithelial development. Previous studies have shown that in hormonally stimulated EEOs, progesterone serves as the main driver of epithelial differentiation, recapitulating *in vivo* patterns [21]. This hormonal stimulation results in increased proportions of glandular and ciliated epithelial cells, alongside a reduction in endometrial epithelial stem cell populations [36]. Within this framework, the differential regulation induced by 10 mM D-lac suggests that this metabolite may enhance progesterone-driven effects, promoting epithelial growth, maturation and differentiation, thereby supporting a receptive state of the endometrium.

In contrast to the modulatory effects of D-lac, exposure to LPS triggers distinct transcriptional responses in EEOs, reflecting the activation of inflammatory pathways. At the lowest concentration tested (2.5 ng/mL), early changes in cytokine-related pathways suggest a shift in the local balance of immune signals, a critical aspect during implantation. Precise regulation of these signals is essential to allow blastocyst acceptance while maintaining defence against microbes [37]. Activation of immune pathways could therefore interfere with the establishment of an optimal receptive state. Given that the LPS used in this study is derived from *E. coli*, this observation is particularly relevant, as colonization by this bacterium has been linked to impaired implantation-related processes [38]. Moreover, activation of pathways related to *E. coli* infection has been reported to correlate with recurrent implantation failure, suggesting that *E. coli*–associated signals may disrupt key molecular mechanisms required for endometrial receptivity [39].

Among the factors contributing to the enrichment of cell development pathways in EEOs exposed to 5 ng/mL LPS, *OLFM4* emerges as a relevant gene. Previous studies have described *OLFM4* as a hormonally regulated marker of endometrial epithelial proliferation, induced by oestrogen and predominantly expressed during the proliferative phase [21]. In our model, *OLFM4* was upregulated in hormonally stimulated EEOs exposed to 5 ng/mL LPS, suggesting that inflammatory signalling interferes with normal hormone-driven epithelial maturation. At this concentration, LPS may therefore sustain a proliferative-like state, impairing the transition to the secretory phase and potentially compromising endometrial receptivity. At the highest LPS concentration tested (10 ng/mL), signalling mediated by the ErbB receptor family, including *ERBB2*, is enriched. This pathway promotes cell proliferation and opposes apoptosis, and its strict regulation is essential to prevent uncontrolled cellular growth. Evidence from mouse models indicates that dysregulation of ErbB signalling can compromise implantation, highlighting its central role in maintaining endometrial receptivity [40].

The analysis of endometrial receptivity markers in hormonally stimulated EEOs exposed to D-lac and LPS provides important insight into how these signals may influence the molecular programs underlying the establishment of a receptive state. Overall, most DEGs in EPC-EEOs were also differentially regulated upon exposure to either D-lac or LPS. However, we observed certain genes that were uniquely differentially regulated in response to the exposure of D-lac or LPS. In the case of D-lac, we observed an upregulation of *DYNLT3*, a gene previously described as an epithelial-specific marker of endometrial receptivity [18]. This upregulation suggests that D-lac may modulate epithelial-specific pathways that contribute to the establishment of a receptive state. This selective induction in EEOs exposed to 2.5 mM D-lac indicates that D-lac may potentially reinforce molecular programs that support the establishment of a receptive endometrial state. In contrast, exposure to LPS was associated with the upregulation of *MAP3K5*, a kinase involved in inflammatory responses, oxidative stress and apoptosis regulation. While *MAP3K5* activation has been described during normal endometrial receptivity, [41] in the context of LPS exposure it likely reflects a cellular response to the inflammatory stimulus rather than a direct enhancement of receptive status. This interpretation is supported by *in vivo* studies in murine models showing that uterine activation of *MAP3K5* by LPS contributes to inflammatory responses that can trigger adverse pregnancy outcomes, such as preterm birth [42].

The three-dimensional organoid model used consists exclusively of epithelial cells, which limits its ability to recapitulate the complex cellular interactions of the endometrium. In this context, the development of multicellular models incorporating stromal, endothelial and immune cells would represent a key strategy to achieve a more accurate representation of the *in vivo* environment. Additionally, our study focused on individual bacteria-derived compounds, whereas the endometrial environment *in vivo* is exposed to a complex mixture of microbial signals that may interact synergistically or antagonistically, potentially influencing epithelial function in ways not captured by our current model.

Together, these results indicate that D-lac and LPS exert hormonally mediated distinct modulatory effects on endometrial receptivity, with D-lac selectively enhancing epithelial programs associated with receptivity, while LPS primarily triggers an inflammatory response. These subtle alterations highlight the capacity of EEOs to integrate hormonal and microbial signals, providing a framework to understand how specific microbial-derived compounds may influence endometrial function.

## Data availability statement

The data presented in the study are deposited in the NCBI SRA Database, BioProject ID PRJNA1415168.

## Acknowledgements

We also extend our deepest gratitude to the clinical collaborators and the participants for their engagement that made this research possible.

## Author’s contributions

S.A., A.S. and A.S.L. jointly supervised this study. L.B.R. initiated and established the cellular models for the endometrium, performed and participated in all experiments, analysed the data, and wrote the manuscript. A.A. and A.P. assisted with the establishment of the experiments, as well as the analysis and interpretation of the results. D.L. contributed to performing the resazurin assays. M.S. was responsible for tissue processing and the collection of endometrial samples. R.M. contributed to the conceptualization and reviewed the manuscript. S.A. contributed to funding and reviewed the manuscript. A.S. provided guidance and support during the experimental work, contributed to funding and reviewed the manuscript. A.S.L. oversaw the study, contributed to funding acquisition, data interpretation and manuscript revision. All authors reviewed and approved the final version of the manuscript.

## SUPPLEMENTARY FIGURES

**Supplementary Figure 1.**
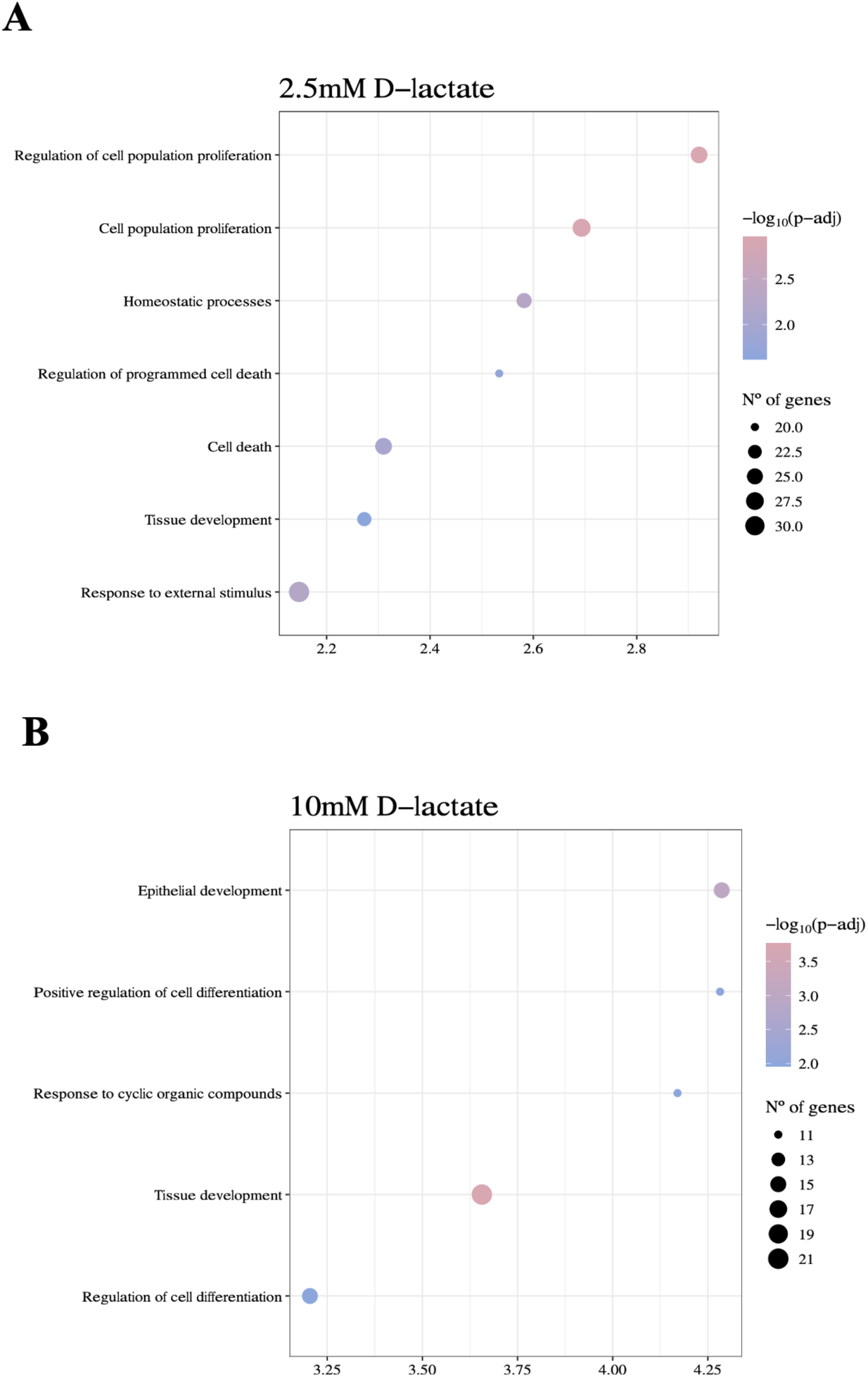
Gene ontology (GO) analysis of upregulated biological pathways in hormonally stimulated endometrial epithelial organoids (EEOs) exposed to D-lactate, compared to EEOs not hormonally stimulated but exposed to D-lactate. Genes differentially expressed exclusively by hormonal action were excluded from this analysis. A) Biologically pathways upregulated in EEOs exposed to 2.5mM D-lactate. B) Biologically pathways positively regulated in EEOs exposed to 10mM D-lactate.

**Supplementary Figure 2.**
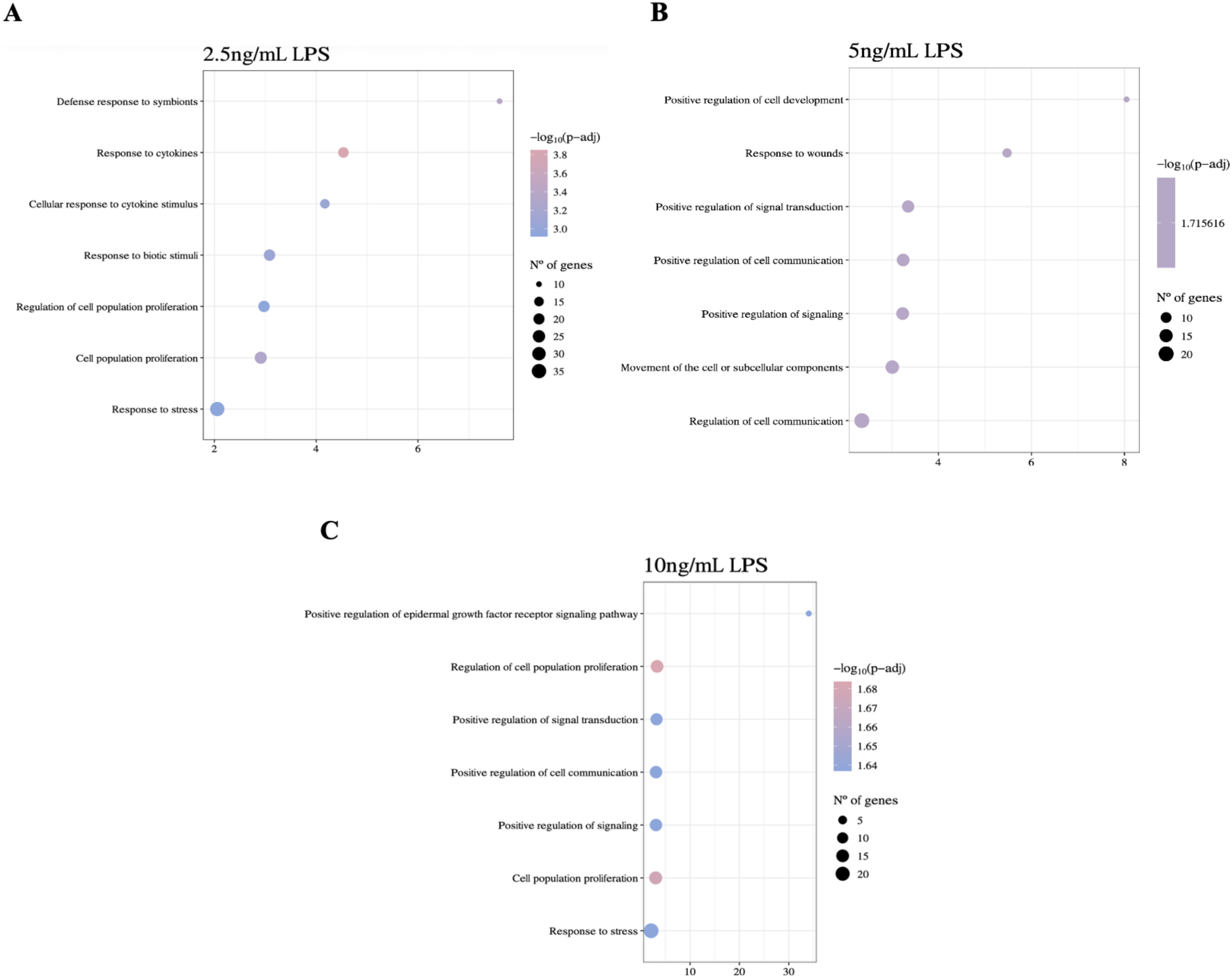
Gene ontology (GO) analysis of positively regulated biological pathways in hormonally stimulated endometrial epithelial organoids (EEOs) exposed to lipopolysaccharide (LPS), compared to EEOs not hormonally stimulated but exposed to LPS. Genes differentially expressed exclusively by hormonal action were excluded from this analysis. A) Biologically pathways positively regulated in EEOs exposed to 2.5ng/mL LPS. B) Biologically pathways positively regulated in EEOs exposed to 5ng/mL LPS. C) Biologically pathways positively regulated in EEOs exposed to 10ng/mL LPS.

